# GraphProt2: A graph neural network-based method for predicting binding sites of RNA-binding proteins

**DOI:** 10.1101/850024

**Authors:** Michael Uhl, Van Dinh Tran, Florian Heyl, Rolf Backofen

## Abstract

CLIP-seq is the state-of-the-art technique to experimentally determine transcriptome-wide binding sites of RNA-binding proteins (RBPs). However, it relies on gene expression which can be highly variable between conditions, and thus cannot provide a complete picture of the RBP binding landscape. This creates a demand for computational methods to predict missing binding sites. Here we present GraphProt2, a computational RBP binding site prediction framework based on graph convolutional neural networks (GCNs). In contrast to current CNN methods, GraphProt2 offers native support for the encoding of base pair information as well as variable length input, providing increased flexibility and the prediction of nucleotide-wise RBP binding profiles. We demonstrate its superior performance compared to GraphProt and two CNN-based methods on single as well as combined CLIP-seq datasets. Conceived as an end-to-end method, GraphProt2 includes all necessary functionalities, from dataset generation over model training to the evaluation of binding preferences and binding site prediction. Various input types and features are supported, accompanied by comprehensive statistics and visualizations to inform the user about datatset characteristics and learned model properties. All this makes GraphProt2 the most versatile and complete RBP binding site prediction method available so far.

## 1 Introduction

RNA-binding proteins (RBPs) regulate many vital steps in the RNA life cycle, such as splicing, transport, stability, and translation [9]. Recent studies suggest a total number of more than 2,000 human RBPs, including 100s of unconventional RBPs, i.e., RBPs lacking known RNA-binding domains [3, 13, 21]. Numerous RBPs have been implicated in diseases like cancer, neurodegeneration, and genetic disorders [4, 8, 31], urging the need to speed up their functional characterization and shed light on their complex cellular interplay.

An important step to understand RBP function is to identify the precise RBP binding locations on regulated RNAs. In this regard, CLIP-seq (cross-linking and immunoprecipitation followed by next generation sequencing) [20] together with its popular modifications PAR-CLIP [12], iCLIP [17], and eCLIP [43] has become the state-of-the-art technique to experimentally determine transcriptome-wide binding sites of RBPs. A CLIP-seq experiment for a specific RBP results in a library of reads bound and protected by the RBP, making it possible to deduce its binding sites by mapping the reads back to the respective reference genome or transcriptome. In practice, computational analysis of CLIP-seq data has to be adapted for each CLIP-seq protocol [40]. Within the analysis, arguably the most critical part is the process of peak calling, i.e., to infer RBP binding sites from the mapped read profiles. Among the many existing peak callers, some popular tools are Piranha [42], CLIPper [22], PEAKachu [2], and PureCLIP [19].

While peak calling is essential to separate authentic binding sites from unspecific interactions and thus reduce the false positive rate, it cannot solve the problem of expression dependency. In order to detect RBP binding sites by CLIP-seq, the target RNA has to be expressed at a certain level in the experiment. Since gene expression naturally varies between conditions, CLIP-seq data cannot be used directly to make condition-independent binding assumptions on a transcriptome-wide scale. Doing so would only increase the false negative rate, i.e., marking all regions not covered by CLIP-seq reads as non-binding, while in fact one cannot tell due to the lack of expression information. Moreover, expression variation is especially high for lncRNAs, an abundant class of ncRNAs gaining more and more attention due to their diverse cellular roles [18]. It is therefore of great importance to infer RBP binding characteristics from CLIP-seq data in order to predict missing binding sites. To give an example, [5] investigated the role of the splicing factor PTBP1 in differential splicing of the tumor suppressor gene ANXA7 in glioblastoma. Despite strong biological evidence for PTBP1 directly binding ANXA7, no binding site was found in a publicly available CLIP-seq dataset for PTBP1. Instead, only a computational analysis was capable to detect and correctly localize the presence of potential binding sites which were then experimentally validated.

Over the years, many approaches to RBP binding site prediction have been presented, from simple sequence motif search to more sophisticated methods incorporating classical machine learning and lately also deep learning. Some popular earlier methods include RNAcontext [15] and GraphProt [23], which can both incorporate RNA sequence and structure information into their predictive models. While RNAcontext utilizes a sequence and structure motif model, GraphProt uses a graph kernel approach, showing improved performance over motif-based techniques. From 2015 on, various deep learning based methods have been proposed, starting with DeepBind [1], which uses sequence information to train a convolutional neural network (CNN). Subsequent methods largely built upon this methodology, using CNNs in combination with recurrent neural networks and additional features such as structure, evolutionary conservation, or region type information [29]. While these methods certainly provide state-of-the-art predictive performance, CNNs by design restrict the input sequences to a fixed length for a given model. Moreover, they cannot encode base pair information, i.e., annotated connections between non-adjacent bases, calling for a more flexible approach that can deal with these limitations, while at the same time supporting additional features.

Here we propose a novel method called GraphProt2 that utilizes a graph convolutional neural network (GCN) to learn binding site properties and predict binding sites. GraphProt2 encodes input sequences as graphs, thus offering native support for base pair information in the form of graph edges. Furthermore, input sequences can be of variable length. This also allows GraphProt2 to calculate binding profiles, by using variable window sizes for computing position-wise binding scores. Binding profiles have proven to be of great practical value, and have been successfully applied in studies such as [7, 27]. In contrast, CNN methods are constrained to fixed-sized inputs. The input size has to be set prior to model training, which is less flexible than directly accepting the variable-size inputs usually defined by peak callers. Moreover, fixed sizes make it more difficult to calculate accurate position-wise profiles, especially near sequence ends. As with CNN methods, GraphProt2 supports additional position-wise features (also termed node attributes), like unpaired probabilities, conservation scores, or region type information. Conceived as an end-to-end method, GraphProt2 includes all the necessary functionalities, from training and test set generation to model training, prediction, and evaluation. Moreover, GraphProt2 is currently the most flexible method with regard to the support of input data types: apart from sequences and genomic regions, it also supports transcript regions, providing automatic feature annotations for all three types. Comprehensive statistics and visualizations are provided as well, in the form of HTML reports, binding profiles, and logos.

Section 2 provides a detailed description of the method, including available modes, features, and data types. In Section 3 we demonstrate GraphProt2’s superior performance, compared to GraphProt and two CNN-based methods (DeepBind and iDeepS), on a set of single CLIP-seq datasets and a combined dataset to learn a generic model. We also show that retrieved sequence logos and binding profiles are in agreement with literature knowledge. Section 4 concludes our findings, sums up existing challenges, and points out possible ways to further improve RBP binding site prediction.

## 2 Methods

### 2.1 The GraphProt2 framework

GraphProt2 utilizes RBP binding sites identified by CLIP-seq and related protocols to train a graph convolutional neural network (GCN) based model, which is then used to predict new binding sites on given input RNA sequences. Figure 1 illustrates the GraphProt2 framework and its general workflow. GraphProt2 requires at least three inputs: a set of RBP binding sites (either in BED or FASTA format), a genomic sequence file (.2bit format), and a genomic annotations file (GTF format). Compared to FASTA, genomes in binary .2bit format occupy less disk space, allow for faster sequence extraction, and also store repeat region information (Section 2.4). Binding sites can be supplied either as sequences, genomic regions, or as transcript regions (GTF file with corresponding transcript annotation required). Additional inputs are available, depending on the binding site input type as well as the selected features (see GitHub repository for a complete documentation). Currently supported features are described in Section 2.4. GraphProt2 is divided into five different program modes: training set generation, prediction set generation, model training, model evaluation, and model prediction (see Supplementary Methods for more details). Depending on the executed mode, various output files are generated. For the dataset generation modes, HTML reports can be output which include detailed statistics and visualizations regarding the positive, negative, or test dataset. This way one can easily compare e.g. the positive input set with the generated negative set and spot possible similarities and differences. Currently there are comparative statistics available on: site lengths, sequence complexity, di-nucleotide distributions, k-mer statistics, target region biotype and overlap statistics, as well as additional statistics and visualizations for each chosen feature. In model evaluation mode, sequence and additional feature logos are output, as well as position-wise scoring profiles for a subset of training sites to illustrate binding preferences (Section 2.6). In model prediction mode, position-wise scoring profiles or whole site predictions are output, and top-scoring sites are extracted from the generated profiles.

**Figure 1.**
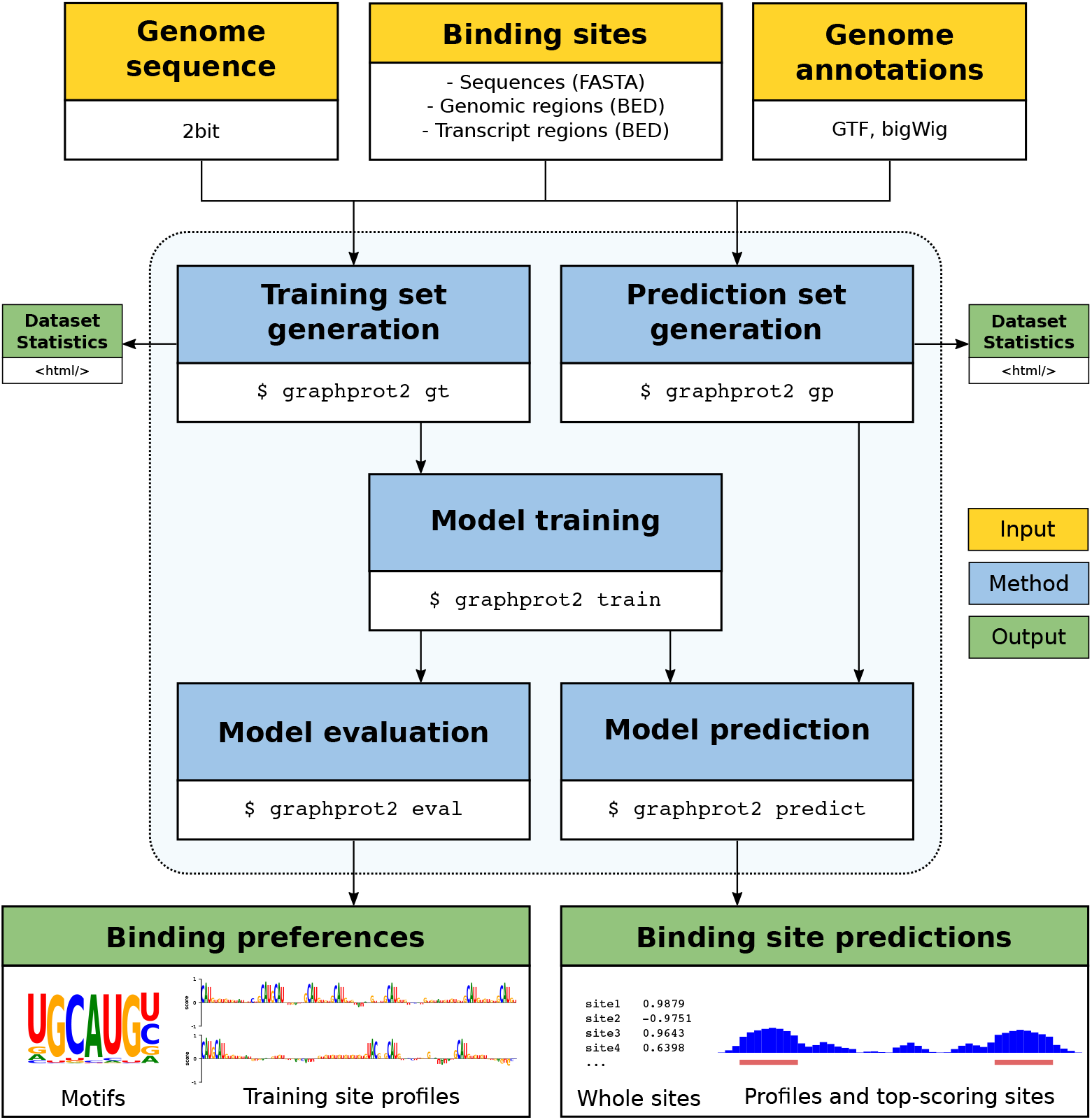
Overview of the GraphProt2 framework. Yellow boxes mark necessary framework inputs, blue boxes the five program modes of GraphProt2, and green boxes the framework outputs. Arrows show the dependencies between inputs, modes, and outputs.

### 2.2 Model architecture

Figure 2 sketches the GraphProt2 model architecture. Given an RBP binding site, its binding site sequence (optionally with secondary structure information) is first transformed into a graph, with nodes corresponding to the nucleotides, and edges to RNA backbone or base pair bonds. Each node gets a node feature or node attributes vector, storing sequence (i.e., its nucleotides as one-hot encoding) and additional feature values (numeric or one-hot encoded). Representations of the graph are then learned by the GCN via several graph convolution layers. Briefly, in each layer the node attributes vector of each node is transformed and updated by aggregating the attribute vectors of adjacent nodes (see Section 2.3 for a formal description). This way, each node aggregates information from an expanding neighborhood as the number of layers increases. Following the convolution layers is a readout phase, to generate fixed-size inputs for the fully connected layers. In the end, the network outputs probabilities for the graph belonging to the positive or negative class, and the maximum probability class determines the predicted class label. For more details on model training and hyperparameters, please refer to the Supplementary Methods or the online documentation.

**Figure 2.**
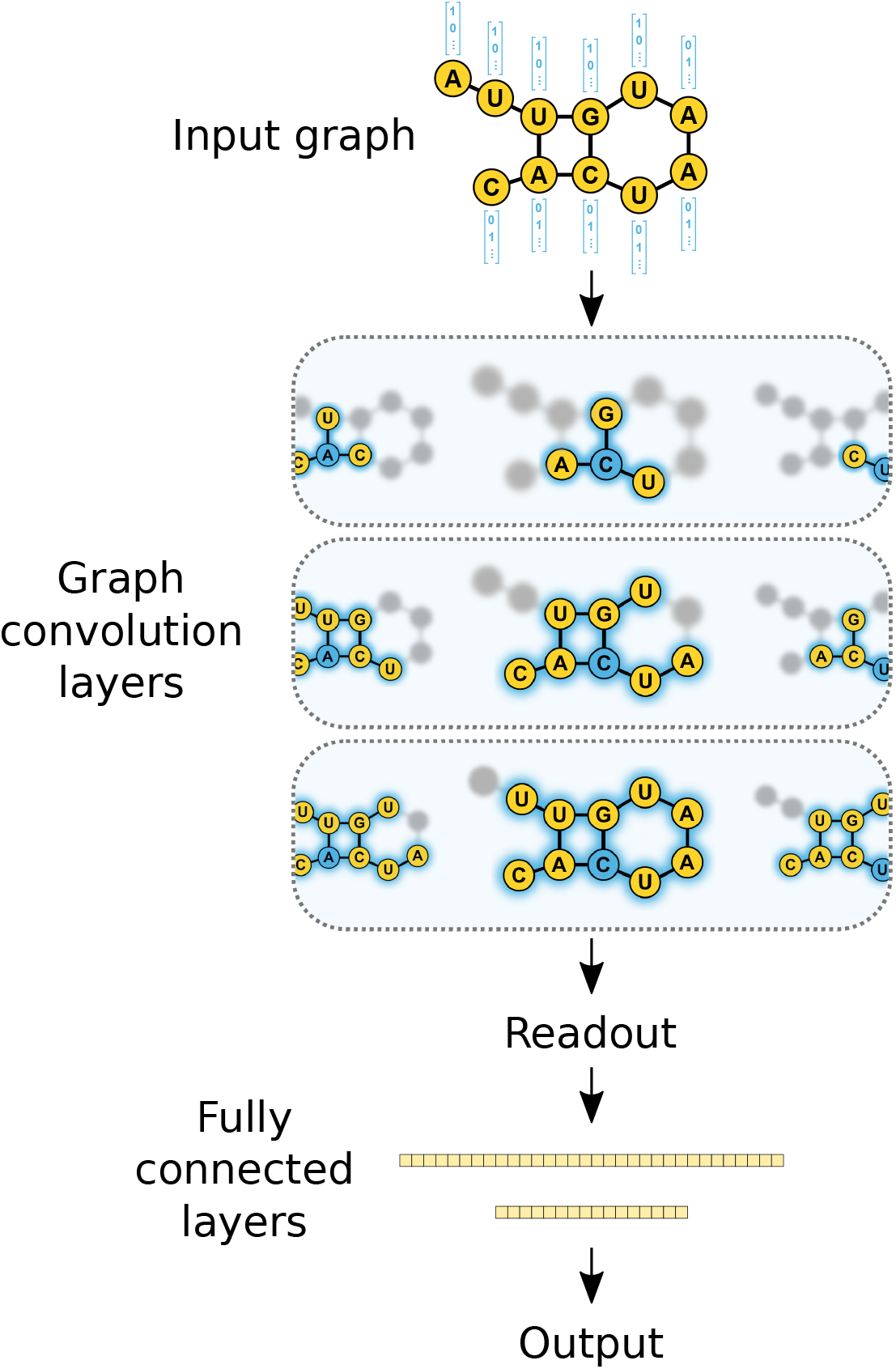
The architecture schematic of Graphprot2 model using three GCN layers.There have been a number of methods proposed to improve this essential form of GCN by, for instance, incorporating graph kernels [24, 25].

### 2.3 Neural network for graphs

In this section we formally describe graph neural networks (GNNs) and provide the definitions necessary to understand the applied graph convolution operations.

#### Notation and definitions

We denote matrices, vectors, and variables with bold uppercase, uppercase, and lowercase letters, respectively. We consider a graph as a tuple *G* = (𝕍,𝔼,**X**), where (𝕍, 𝔼 are the sets of nodes and edges. **X** ∈ ℝ^*n×d*^ is a node attribute matrix, where each row **X**_*i*_ is a real-valued vector of size *d* associated to node *v*_*i*_ of the graph. We define the adjacency matrix **A** ∈ ℝ^*n×n*^ as *a*_*ij*_ = 1 *⇐⇒* (*i, j*) ∈, 𝔼,
and 0 otherwise. **D** is the degree matrix, where *d*_*ij*_ = Σ*_j_ a_ij_* if *i* = *j*, and 0 otherwise.

The first concepts of GNNs were described in [32, 34]. Based on these concepts, many methods have been proposed later on [16, 25, 48]. For an overview of existing GNN methods, we recommend the following recent survey [46]. GraphProt2 employs a GCN, which is a special kind of GNN that uses graph convolutions. A general GCN architecture includes the three main components: graph convolutions, a readout phase, and fully connected layers. In the following, we briefly describe the first two components.

#### Graph convolutions

A graph convolution has its architecture defined following the graph topology, in which nodes are considered as neurons and edges as links in the network. Each node is assigned a state and the graph convolutions aim at iteratively updating each node state over time. Different policies used to update node states define distinct graph convolutions [16, 39, 45, 48]. Generally, graph convolutions employ the current state of a node together with the accumulation over its neighbouring states, within a pre-defined number of hops, to update the node state.

Given a graph *G* with **A, X** as its adjacency and attribute matrix, we use the following graph convolution as described in [16]:

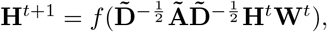

where **H**^**0**^ = **X** and *L* is the number of convolution layers with *t* = 0 … (*L* 1). **H**^*t*^ is the state matrix or convolution output at time *t*, **Ã** = **A** + **I** with **I** being the identity matrix, and 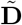 is the degree matrix corresponding to **Ã**. **W**^*t*^ is the weight matrix containing the trainable convolution parameters at time *t*, and *f* is the element-wise non-linear activation function.

#### Readout phase

Following the graph convolutions is a readout phase, in which the variable-size convolution outputs are converted into fixed-size inputs for the fully connected layers. In particular, graph node representations are taken from all convolution outputs of a graph and converted into a vector of fixed size, consistent over all graphs. A number of readout methods have been proposed in [26, 48]. For GraphProt2, we first take the maximum, minimum, and median of each node feature over all node feature vectors of a graph. We then use their concatenation to get a fixed-size feature vector representing the graph.

### 2.4 Supported features

GraphProt2 currently supports the following position-wise features which can be utilized for training and prediction in addition to the sequence feature: secondary structure information (base pairs and structural element probabilities), conservation scores (phastCons and phyloP), exon-intron annotation, transcript region annotation, and repeat region annotation. Table 1 lists the features available for each binding site input type.

**Table 1.** GraphProt2’s three supported input types (sequences, genomic regions, transcript regions) and the features available for them.

| Feature | Sequences | Genomic regions | Transcript regions |
| --- | --- | --- | --- |
| structure | YES | YES | YES |
| conservation scores | NO | YES | YES |
| exon-intron annotation | NO | YES | NO |
| transcript region annotation | NO | YES | YES |
| repeat region annotation | NO | YES | YES |

#### Secondary structure information

GraphProt2 can include two kinds of structure information for a given RNA sequence: 1) base pairs and 2) unpaired probabilities for different loop contexts (probabilities for the nucleotide being paired or inside external, hairpin, internal or multi loops). Both are calculated using the ViennaRNA Python 3 API (ViennaRNA 2.4.14) and RNAplfold with its sliding window approach, with user-definable parameters (by default these are window size = 150, maximum base pair span length = 100, and probabilities for regions of length u = 3). The base pairs with a probability≥ a set threshold (default = 0.5) are then added to the sequence graph as edges between the nodes that represent the end points of the base pair, and the unpaired probability values are added to the node feature vectors. Alternatively, the user can also provide custom base pair information, e.g. derived from experimental data.

#### Conservation scores

GraphProt2 supports two scores measuring evolutionary conservation (phastCons and phyloP). Conservation scores were downloaded from the UCSC website, using the phastCons and phyloP scores generated from multiple sequence alignments of 99 vertebrate genomes to the human genome (hg38, phastCons100way and phyloP100way datasets). GraphProt2 accepts scores in .bigWig format. To assign conservation scores to transcript regions, transcript regions are first mapped to the genome, using the provided GTF file.

#### Exon-intron annotation

Exon-intron annotation in the form of one-hot encoded exon and intron labels can also be added to the node feature vectors. Labels are assigned to each binding site position by taking a set of genomic exon regions and overlapping it with the genomic binding sites using bedtools (2.29.0). To unambiguously assign labels, GraphProt2 by default uses the most prominent isoform for each gene. The most prominent isoform for each gene gets selected through hierarchical filtering of the transcript information present in the input GTF file (for the benchmark results in Section 3 we used Ensembl Genes 97, GRCh38.p12): given that the transcript is part of the GENCODE basic gene set, GraphProt2 selects transcripts based on their transcript support level (highest priority), and by transcript length (longer isoform preferred). The extracted isoform exons are then used for region type assignment. Note that this feature is only available for genomic regions, as it is not informative for transcript regions, which would contain only exon labels. Optionally, a user-defined isoform list can be supplied, substituting the list of most prominent isoforms for annotation. Regions not overlapping with introns or exons can also be labelled separately (instead of labelled as intron).

#### Transcript region annotation

Similarly to the exon-intron annotation, binding regions can be labelled based on their overlap with transcript regions. Labels are assigned based on UTR or CDS region overlap (5’UTR, CDS, 3’UTR, None), by taking the isoform information in the input GTF file. Again the list of most prominent isoforms is used for annotation, or alternatively a list of user-defined isoforms. Additional annotation options include start and stop codon or transcript and exon border labelling.

#### Repeat region annotation

Repeat region annotation can also be added to the binding regions, analogously to other region type annotations. This information is derived directly from the genomic sequence file (in .2bit format, from the UCSC website), where repeat regions identified by RepeatMasker and Tandem Repeats Finder are stored in lowercase letters.

### 2.5 Tool benchmark

#### Dataset construction

To construct the benchmark sets we extracted eCLIP data out of two cell lines (HepG2, K562) from the ENCODE project website [33] (November 2018 release). We directly used the genomic binding regions (genome assembly GRCh38) identified by ENCODE’s in-house peak caller CLIPper, which are available in BED format for each RBP and each replicate, often for both cell lines (thus 4 replicate BED files per RBP). For the single model benchmark set, binding sites were further filtered by their log2 fold change (FC) to obtain*∼*6,000 to 10,000 binding regions for each replicate. We next removed sites with length *>* 0.75 percentile length and selected for each RBP the replicate set that contained the most regions, centered the sites, and extended them to make all sites equal length. We chose a binding site length of 101 nt (50 nt extension up- and downstream of center position) and randomly selected 30 sets.

For the generic model benchmark set, we first merged both replicates of each RBP cell type combination, keeping only the sites with the highest FC in case of overlapping sites. After filtering (FC = 1), centering, and extending sites to 61 nt, we clustered the RBP cell type combinations (120 for K562, 104 for HepG2) by their binding site 3-mer content. We selected k=6 (maximum silhouette score), and selected 2 to 5 sets from each cluster, resulting in 20 datasets. After filtering (FC = 3) and randomly choosing 2,000 sites for each set, we again only kept the top FC sites in case of overlaps, and normalized the site lengths to 101 nt. This resulted in a set of 38,978 RBP binding sites from 20 different RBPs.

To generate the negative sets for both single and generic sets, we randomly selected sites based on two criteria: 1) their location on genes covered by eCLIP peak regions and 2) no overlap with any eCLIP peak regions from the experiment. The same number of random negative and positive instances was used throughout the benchmarks.

#### Tool setup and performance measurement

GraphProt2 and iDeepS were benchmarked with default parameters. For DeepBind, we used the implementation available in DeepRAM [38], including hyperparameter optimization. For GraphProt, hyperparameters were optimized for each model before cross validation, using a separate optimization set of 500 positive and 500 negative examples. The accuracy measure, i.e., the proportion of correctly classified instances, was used in combination with 10-fold cross validation to estimate and compare single model generalization performances. Standard deviation of the 10 measured accuracies is reported for each model, together with the average accuracy (DeepBind, iDeepS, GraphProt2). Note that we could not provide standard deviations for GraphProt, as it does not ouput the single fold accuracies. For generic models with *n* RBPs, the sites of each RBP were selected once for testing, while the remaining *n−*1 RBP sites were used for training, in total generating *n* different accuracy measures.

### 2.6 Visualization

To visualize learned binding properties, GraphProt2 can output sequence and additional feature logos of various lengths, as well as position-wise scoring profiles for a user-defined subset of training sites (see Figure 4 and Supplementary Figure 1 for visualization examples). The support of variable-sized inputs (i.e., graphs of variable size) allows GraphProt2 to calculate position-wise scoring profiles using a sliding window approach. Each window corresponds to a subgraph, which is scored by the model, and the score is assigned to the center (backbone) position of the graph. To generate a logo, GraphProt2 extracts top-scoring positions from a specified number of top scoring positive site profiles, and extends them to a defined logo length. The extracted subsequences are then converted into a weight matrix and plotted with Logomaker [37].

**Figure 3.**
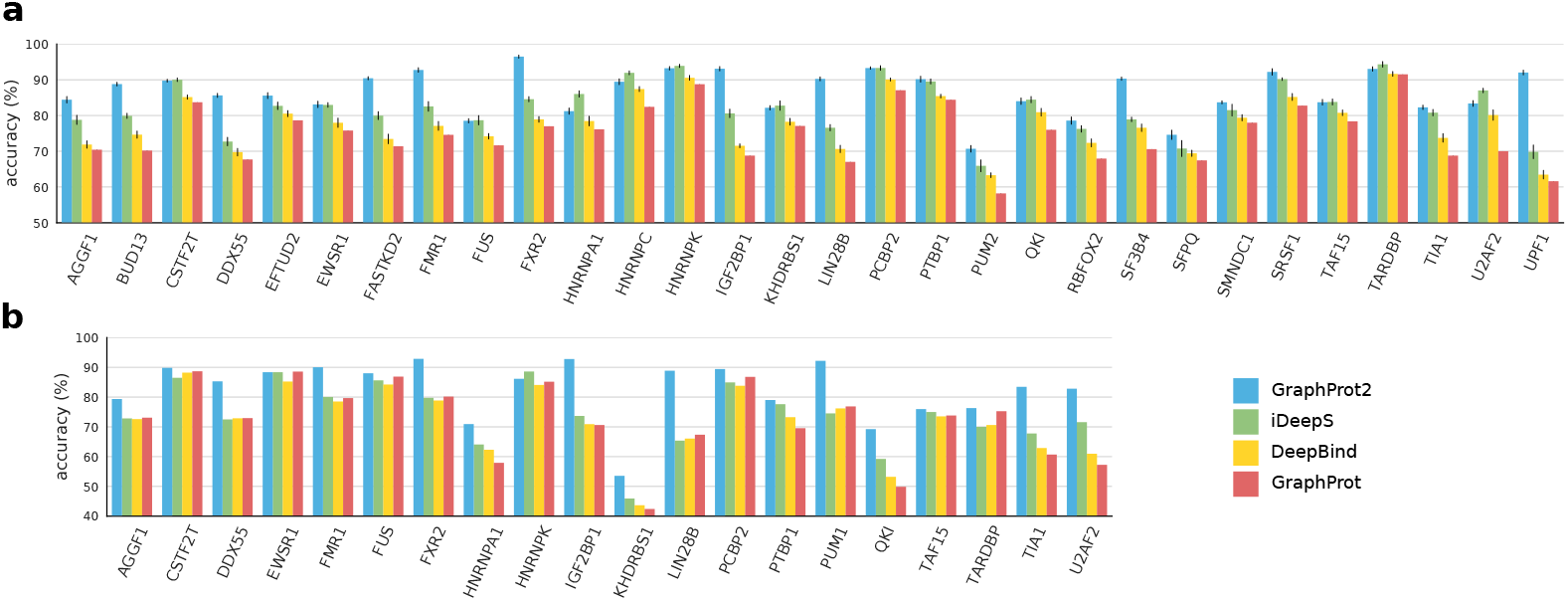
Model benchmark results for GraphProt2, iDeepS, DeepBind, and GraphProt. (a) Single model results over 30 individual RBP eCLIP sets. We plot average accuracies obtained by 10-fold cross validation together with standard deviations (apart from GraphProt). (b) Generic model results over a combined eCLIP set containing sites from 20 different RBPs. In each round a model was trained on 19 RBP sets and tested on the remaining RBP set. We plot test accuracies for each round, together with the name of the test set RBP.

**Figure 4.**
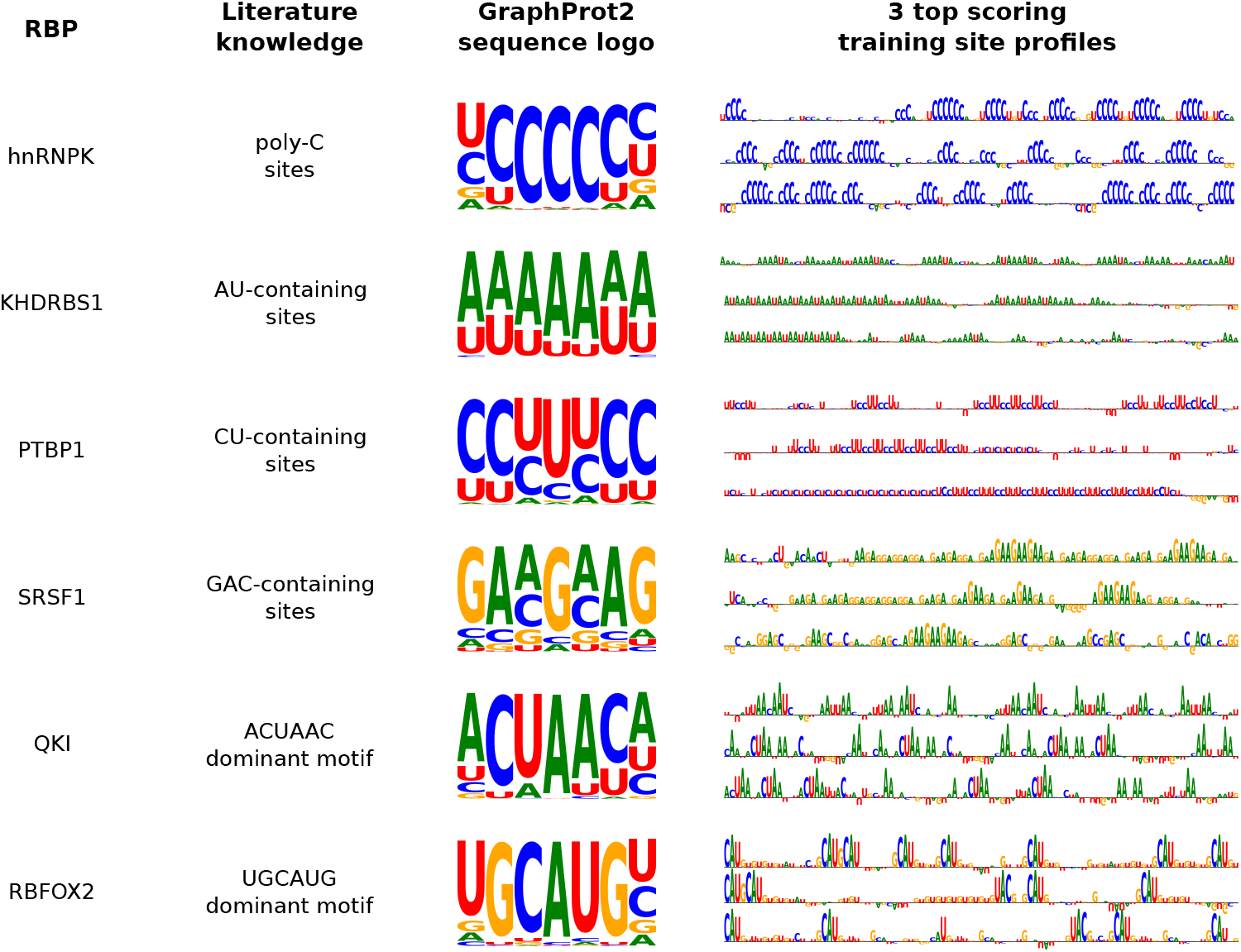
Comparison of GraphProt2 sequence logos and profiles with known RBP binding preferences. Literature knowledge was obtained from the ATtRACT database [11]. All models were trained using only the sequence feature for easier interpretation. Logos were generated by extracting the top scoring position for each of the top 500 scoring profiles, and an extension of 3 nt on each side to create 500 subsequences of length 7. Logo character heights thus correspond to character frequencies at each of the 7 positions (centered on the top scoring position). Window sizes of 5 and 7 were used to calculate the profiles, where the character height corresponds to the window score (i.e., the positive class probability of the subgraph).

### 2.7 Tool requirements and availability

GraphProt2 is Python-based and uses the PyTorch framework [30] in combination with PyTorch Geometric [6] to enable deep learning for graphs. It is possible to run GraphProt2 completely on CPU, but recommended to use an Nvidia GPU (CUDA 10 required) to speed up computations. Detailed requirements, installation instructions, as well as the paper supplements can be found on the tool GitHub site: https://github.com/BackofenLab/GraphProt2.

## 3 Results and discussion

In the following, we demonstrate GraphProt2’s superior performance based on two benchmarks, one over 30 single eCLIP RBP datasets, and the other over a combined set containing data from 20 RBPs. We compared GraphProt2 with GraphProt, a graph kernel approach that uses an SVM classifier, DeepBind, the first CNN-based method, and iDeepS [28], a CNN-based method that also incorporates a long short-term memory (LSTM) architecture. For iDeepS, we trained models using both sequence and structure information. For GraphProt, we know from former studies that sequence models usually perform similar to structure models, while taking considerably less time for training. We therefore chose to train sequence models for the comparison. For GraphProt2, we found that incorporating structure into the graphs also did not significantly change the performance for the described benchmark sets. We therefore restricted the node features to sequence, conservation scores, and exon-intron annotation for the two benchmarks. We did not include transcript region annotation for the benchmark, since accurate (i.e., condition-specific) CDS and UTR region annotation requires additional analysis tools for the detection of expressed isoforms. Repeat region annotation was also not included, since the eCLIP pipeline that produced the binding sites used for training only considers uniquely mapped reads. However, a recent pipeline update [44] now also allows mapping to certain repeat elements, and has already lead to the discovery of many new RBP binding sites overlapping with these elements. Repeat region annotation could thus become an informative feature once these datasets are available.

### 3.1 GraphProt2 performs superior over single models

Figure 3a shows the 10-fold cross validation results over 30 single eCLIP sets for GraphProt, iDeepS, DeepBind, and GraphProt2. GraphProt2 achieves the highest total average accuracy (86.43%), followed by iDeeps (82.24%), DeepBind (77.63%), and GraphProt (74.66%) (see Supplementary Table 2 for values). This substantial increase in accuracy clearly demonstrates the power of deep learning methods compared to earlier state-of-the-art methods like GraphProt. In all 30 cases, GraphProt is outperformed by the deep learning methods (except of a few ties with DeepBind). GraphProt2 shows better performance in 17 cases, while 9 cases are ties between GraphProt2 and iDeepS, and in 5 cases iDeepS performs best.

Looking closer at the 17 cases, we can sometimes see drastic improvements (*>* 10%), e.g. for DDX55, IGF2BP1, LIN28B, or UPF1. Given that all three methods use the same sequences for training, we found that these large performance increases can be mainly attributed to either one or two of the additional features (conservation or exon-intron annotation). One could argue that the additional features mainly capture common biases between positive and negative sets. However, as we do not observe these drastic improvements for the majority of cases, we can assume that they indeed represent RBP specific properties, and thus are informative and valid features for prediction. As for the 5 cases in which iDeepS performs best, the performance increase is less pronounced, with a maximum of ∼4%. Besides architectural differences and the added structure information, the increase could also be due to the incorporated LSTM, which in theory should be able to better identify recurring patterns, but in practice we could not measure its contribution as it cannot be disabled.

It is known that RNA structure can be important for RBP binding [14, 36]. Since structure features did not significantly improve performance in GraphProt2, it could be argued that deep learning methods are powerful enough to detect discriminative structure information directly from the sequence data, whereas in earlier methods like GraphProt, predicted structures were still shown to boost performance for a small number of RBP datasets. Other reasons for the ineffectiveness of structure features could be that they are after all computationally predicted, and that CLIP-seq protocols tend to recover less structured binding sites because crosslinking of double-stranded regions is less efficient. Also, RNA structure itself is highly dynamic and affected by a multitude of interacting RBPs in the cell, which is currently not modelled by any prediction method. In addition, it is not clear whether the structure encoding chosen for a model is optimal for the task. Approaches like adding experimental structure probing data to support predictions, or CLIP-seq protocols that better capture structured binding sites might be a way to reevaluate the importance of structure in RBP binding site prediction. For example, hiCLIP [35] can identify double-stranded regions bound by an RBP, and irCLIP [47] can potentially resolve RNA secondary structures to increase crosslinking efficiency for structured sites. However, these protocols have not yet been widely applied.

### 3.2 GraphProt2 generic model outperforms other methods

In contrast to single models, generic models are trained by combining several RBP datasets, to learn binding properties shared between different RBPs. Such a model can be used to predict binding sites for a studied RBP in cases where there is no CLIP-seq data available, or in general to estimate the likelihood of RBP binding for a given sequence. Figure 3b presents the generic model results over a combined eCLIP set, consisting of sites from 20 different RBPs. As with the single models, GraphProt2 obtains the highest average accuracy (82.49%), followed by iDeepS (74.19%), GraphProt (72.17%), and DeepBind (72.09%) (see Supplementary Table 3 for values). Out of the 20 test comparisons, GraphProt2 achieves the highest accuracy in 13 cases. Furthermore, there are 6 tie cases between GraphProt2 and the other methods and one case where iDeepS performs best. Looking at the benchmark results, we can see that GraphProt2 increases its average accuracy lead from ~ 4% for single models to ~ 8%, while GraphProt is closer to iDeepS and practically even with DeepBind.

As described, the 20 RBPs were chosen based on k-means clustering of the 3-mer contents of their binding site sequences, in order to create a training set that contains a diverse collection of binding sites from RBPs with different binding preferences. This way we wanted to mitigate biases that would be introduced by random sampling of RBPs, assuming that many RBPs share similar binding preferences. Indeed, we observe that not all RBPs work well as test sets. There are particularly weak performing RBP sets over all three methods, such as for KHDRBS1, QKI, or HNRNPA1. Since we do not experience these drops with their single models, we can assume that these test sets indeed contain useful information, although the information does not seem to be common to most other RBPs in the training set. These varying performances also speak against a strong protocol-specific bias inherent to eCLIP data, which, if present, should result in more similar performances.

Apart from KHDRBS1, GraphProt2 often performs notably better on sets that show low performances (∼ 70% or less) for iDeepS, DeepBind, and GraphProt (DDX55, IGF2BP1, LIN28B, TIA1, and U2AF2). As with the single models, this effect can be attributed to the added conservation and region type information. One could argue that RBP binding sites are naturally biased towards conserved regions or specific region types, which leads to better accuracy scores. On the other hand, these biases also display generic properties of RBP binding sites, and thus are indeed valid features to use. This is especially true when the goal is to train a generic model, which first and foremost needs to discriminate between common RBP binding and non-binding sites.

### 3.3 GraphProt2 models capture known binding preferences

As deep learning models are complex by design and thus hard to interpret, the development of visualizations that help to explain what is learned by a model is an important and active area of research. Since there are currently no readily available visualizations for pyTorch Geometric models, we implemented our own visualization, by taking advantage of GraphProt2’s variable-sized input support (see Section 2.6 for details). To compare GraphProt2 sequence logos and profiles with known RBP binding preferences from the literature, we trained sequence models on six different RBP datasets. Figure 4 shows the obtained sequence logo and known preferences (based on RBP motifs listed in the ATtRACT database [11]), as well as three top scoring profiles for each RBP. As we can see, the logos clearly capture the literature preferences, both for RBPs without a single dominant motif (hnRNPK, KHDRBS1, PTBP1, SRSF1), and for RBPs with strong individual motifs (QKI, RBFOX2). Profiles also agree with literature preferences, although it has to be noted that these visualizations can only provide a simplified view on what the model has actually learned, and what it actually uses to distinguish between positive and negative sites. As visualization solutions for pyTorch geometric are currently in development, e.g. to visualize the importance of single nodes and edges, we are confident to introduce additional visualizations in the near future.

## 4 Conclusion

In this work we presented GraphProt2, a highly versatile deep learning-based RBP binding site prediction method which supports variable length inputs, various input types, and additional features to achieve state-of-the-art predictive performance. Moreover, all functionalities necessary to run RBP binding site predictions are included, making GraphProt2 the most complete available method so far. Compared to popular CNN methods, the ability to work with variable length inputs makes GraphProt2 more flexible and allows for position-wise profile predictions with variable window sizes.

Base pair annotation as well as nucleotide-wise loop context probabilities are also supported, although our results did not show any performance gains for the constructed eCLIP benchmark datasets when adding these features. As discussed in Section 3, there are many possible reasons for this lack of improvement, and it is not clear at this point whether it is worth the extra effort to incorporate structure information into future deep learning methods. Condition-specific experimental structure data could help to reassess the importance of structure, and we would like to evaluate this with GraphProt2, once datasets become available.

To further improve RBP binding site predictions, there are many possible ways, such as new or adapted methods, better visualizations, or improvements in dataset quality. To give some examples, multi-class models trained on multiple RBP datasets to learn common and unique RBP binding patterns can yield a deeper understanding of RBP binding, beyond of what is possible with single RBP models [10]. Regarding visualizations, we will undoubtedly see improvements for GCNs in the near future, due to ongoing development and a generally high interest in deep learning libraries. For example, we would like to visualize single node and edge importances at one point, and if possible also show the importance of individual features. As for the improvement of dataset quality, one drawback current peak callers have is that they do not take into account information from splicing events for binding site identification. This strategy of course is far from optimal, especially for RBPs that bind predominantly to spliced RNA. Indeed, a recent study has shown that it frequently leads to falsely called peaks at exon borders [41]. Moreover, it was shown that context choice influences RBP binding site predictions. It therefore can be assumed that context-sensitive peak calling in combination with transcript region supporting prediction methods will lead to more accurate binding models in the near future.

## Acknowledgments

We thank Daniel Maticzka for his initial work on the method and Martin Raden for his invaluable suggestions on the topic.

## References

1. B. Alipanahi, A. Delong, M. T. Weirauch, and B. J. Frey. Predicting the sequence specificities of dna-and rna-binding proteins by deep learning. Nature biotechnology, 33(8):831, 2015.

2. T. Bischler, D. Maticzka, K. U. Förstner, and P. R. Wright. PEAKachu. https://github.com/tbischler/PEAKachu.

3. K. W. Brannan, W. Jin, S. C. Huelga, C. A. Banks, J. M. Gilmore, L. Florens, M. P. Washburn, E. L. Van Nostrand, G. A. Pratt, M. K. Schwinn, et al. Sonar discovers rna-binding proteins from analysis of large-scale protein-protein interactomes. Molecular cell, 64(2):282–293, 2016.

4. E. G. Conlon and J. L. Manley. Rna-binding proteins in neurodegeneration: mechanisms in aggregate. Genes & development, 31(15):1509–1528, 2017.

5. R. Ferrarese, G. R. Harsh, A. K. Yadav, E. Bug, D. Maticzka, W. Reichardt, S. M. Dombrowski, T. E. Miller, A. P. Masilamani, F. Dai, et al. Lineage-specific splicing of a brain-enriched alternative exon promotes glioblastoma progression. The Journal of clinical investigation, 124(7):2861–2876, 2014.

6. M. Fey and J. E. Lenssen. Fast graph representation learning with PyTorch Geometric. In ICLR Workshop on Representation Learning on Graphs and Manifolds, 2019.

7. A. R. Gawronski, M. Uhl, Y. Zhang, Y.-Y. Lin, Y. S. Niknafs, V. R. Ramnarine, R. Malik, F. Feng, A. M. Chinnaiyan, C. C. Collins, et al. Mechrna: prediction of lncrna mechanisms from rna–rna and rna–protein interactions. Bioinformatics, 34(18):3101–3110, 2018.

8. S. Gerstberger, M. Hafner, M. Ascano, and T. Tuschl. Evolutionary conservation and expression of human rna-binding proteins and their role in human genetic disease. In Systems biology of RNA binding proteins, pages 1–55. Springer, 2014.

9. S. Gerstberger, M. Hafner, and T. Tuschl. A census of human rna-binding proteins. Nature Reviews Genetics, 15(12):829, 2014.

10. M. Ghanbari and U. Ohler. Deep neural networks for interpreting rna-binding protein target preferences. Genome research, 30(2):214–226, 2020.

11. G. Giudice, F. Sánchez-Cabo, C. Torroja, and E. Lara-Pezzi. Attract—a database of rna-binding proteins and associated motifs. Database, 2016, 2016.

12. M. Hafner, M. Landthaler, L. Burger, M. Khorshid, J. Hausser, P. Berninger, A. Rothballer, M. Ascano Jr, A.-C. Jungkamp, M. Munschauer, et al. Transcriptome-wide identification of rna-binding protein and microrna target sites by par-clip. Cell, 141(1):129–141, 2010.

13. M. W. Hentze, A. Castello, T. Schwarzl, and T. Preiss. A brave new world of rna-binding proteins. Nature Reviews Molecular Cell Biology, 19(5):327, 2018.

14. E. Jankowsky and M. E. Harris. Specificity and nonspecificity in rna–protein interactions. Nature reviews Molecular cell biology, 16(9):533–544, 2015.

15. H. Kazan, D. Ray, E. T. Chan, T. R. Hughes, and Q. Morris. Rnacontext: a new method for learning the sequence and structure binding preferences of rna-binding proteins. PLoS computational biology, 6(7):e1000832, 2010.

16. T. N. Kipf and M. Welling. Semi-supervised classification with graph convolutional networks. arXiv preprint arXiv:1609.02907, 2016.

17. J. König, K. Zarnack, G. Rot, T. Curk, M. Kayikci, B. Zupan, D. J. Turner, N. M. Luscombe, and J. Ule. iclip reveals the function of hnrnp particles in splicing at individual nucleotide resolution. Nature structural & molecular biology, 17(7):909, 2010.

18. A. E. Kornienko, C. P. Dotter, P. M. Guenzl, H. Gisslinger, B. Gisslinger, C. Cleary, R. Kralovics, F. M. Pauler, and D. P. Barlow. Long non-coding rnas display higher natural expression variation than protein-coding genes in healthy humans. Genome biology, 17(1):14, 2016.

19. S. Krakau, H. Richard, and A. Marsico. PureCLIP: capturing target-specific protein–RNA interaction footprints from single-nucleotide CLIP-seq data. Genome biology, 18(1):240, 2017.

20. D. D. Licatalosi, A. Mele, J. J. Fak, J. Ule, M. Kayikci, S. W. Chi, T. A. Clark, A. C. Schweitzer, J. E. Blume, X. Wang, et al. Hits-clip yields genome-wide insights into brain alternative rna processing. Nature, 456(7221):464, 2008.

21. L. Liu, T. Li, G. Song, Q. He, Y. Yin, J. Y. Lu, X. Bi, K. Wang, S. Luo, Y.-S. Chen, et al. Insight into novel rna-binding activities via large-scale analysis of lncrna-bound proteome and idh1-bound transcriptome. Nucleic acids research, 47(5):2244–2262, 2019.

22. M. T. Lovci, D. Ghanem, H. Marr, J. Arnold, S. Gee, M. Parra, T. Y. Liang, T. J. Stark, L. T. Gehman, S. Hoon, et al. Rbfox proteins regulate alternative mRNA splicing through evolutionarily conserved RNA bridges. Nature structural & molecular biology, 20:1434, 2013.

23. D. Maticzka, S. J. Lange, F. Costa, and R. Backofen. Graphprot: modeling binding preferences of rna-binding proteins. Genome biology, 15(1):R17, 2014.

24. N. Navarin, D. Tran, and A. Sperduti. Learning kernel-based embeddings in graph neural networks. In European conference on artificial intelligence, 2020.

25. N. Navarin, D. V. Tran, and A. Sperduti. Pre-training graph neural networks with kernels. arXiv preprint arXiv:1811.06930, 2018.

26. N. Navarin, D. Van Tran, and A. Sperduti. Universal readout for graph convolutional neural networks. In 2019 International Joint Conference on Neural Networks (IJCNN), pages 1–7. IEEE, 2019.

27. Y. S. Niknafs, S. Han, T. Ma, C. Speers, C. Zhang, K. Wilder-Romans, M. K. Iyer, S. Pitchiaya, R. Malik, Y. Hosono, et al. The lncrna landscape of breast cancer reveals a role for dscam-as1 in breast cancer progression. Nature communications, 7:12791, 2016.

28. X. Pan, P. Rijnbeek, J. Yan, and H.-B. Shen. Prediction of rna-protein sequence and structure binding preferences using deep convolutional and recurrent neural networks. BMC genomics, 19(1):511, 2018.

29. X. Pan, Y. Yang, C.-Q. Xia, A. H. Mirza, and H.-B. Shen. Recent methodology progress of deep learning for rna–protein interaction prediction. Wiley Interdisciplinary Reviews: RNA, page e1544, 2019.

30. A. Paszke, S. Gross, S. Chintala, G. Chanan, E. Yang, Z. DeVito, Z. Lin, A. Desmaison, L. Antiga, and A. Lerer. Automatic differentiation in PyTorch. In NIPS Autodiff Workshop, 2017.

31. B. Pereira, M. Billaud, and R. Almeida. Rna-binding proteins in cancer: old players and new actors. Trends in cancer, 3(7):506–528, 2017.

32. F. Scarselli, M. Gori, A. C. Tsoi, M. Hagenbuchner, and G. Monfardini. The graph neural network model. IEEE Transactions on Neural Networks, 20(1):61–80, 2009.

33. C. A. Sloan, E. T. Chan, J. M. Davidson, V. S. Malladi, J. S. Strattan, B. C. Hitz, I. Gabdank, A. K. Narayanan, M. Ho, B. T. Lee, et al. Encode data at the encode portal. Nucleic acids research, 44(D1):D726–D732, 2015.

34. A. Sperduti and A. Starita. Supervised neural networks for the classification of structures. IEEE Transactions on Neural Networks, 8(3):714–735, 1997.

35. Y. Sugimoto, A. Vigilante, E. Darbo, A. Zirra, C. Militti, A. D’Ambrogio, N. M. Luscombe, and J. Ule. hiclip reveals the in vivo atlas of mrna secondary structures recognized by staufen 1. Nature, 519(7544):491, 2015.

36. J. M. Taliaferro, N. J. Lambert, P. H. Sudmant, D. Dominguez, J. J. Merkin, M. S. Alexis, C. A. Bazile, and C. B. Burge. Rna sequence context effects measured in vitro predict in vivo protein binding and regulation. Molecular cell, 64(2):294–306, 2016.

37. A. Tareen and J. B. Kinney. Logomaker: beautiful sequence logos in python. Bioinformatics, 36(7):2272–2274, 2020.

38. A. Trabelsi, M. Chaabane, and A. Ben-Hur. Comprehensive evaluation of deep learning architectures for prediction of dna/rna sequence binding specificities. Bioinformatics, 35(14):i269–i277, 2019.

39. D. V. Tran, N. Navarin, and A. Sperduti. On filter size in graph convolutional networks. In 2018 IEEE Symposium Series on Computational Intelligence (SSCI), pages 1534–1541. IEEE, 2018.

40. M. Uhl, T. Houwaart, G. Corrado, P. R. Wright, and R. Backofen. Computational analysis of clip-seq data. Methods, 118:60–72, 2017.

41. M. Uhl, D. Van Tran, and R. Backofen. The importance of incorporating transcript information in clip-seq data analysis. 2020.

42. P. J. Uren, E. Bahrami-Samani, S. C. Burns, M. Qiao, F. V. Karginov, E. Hodges, G. J. Hannon, J. R. Sanford, L. O. Penalva, and A. D. Smith. Site identification in high-throughput rna–protein interaction data. Bioinformatics, 28(23):3013–3020, 2012.

43. E. L. Van Nostrand, G. A. Pratt, A. A. Shishkin, C. Gelboin-Burkhart, M. Y. Fang, B. Sundararaman, S. M. Blue, T. B. Nguyen, C. Surka, K. Elkins, et al. Robust transcriptome-wide discovery of rna-binding protein binding sites with enhanced clip (eclip). Nature methods, 13(6):508, 2016.

44. E. L. Van Nostrand, G. A. Pratt, B. A. Yee, E. C. Wheeler, S. M. Blue, J. Mueller, S. S. Park, K. E. Garcia, C. Gelboin-Burkhart, T. B. Nguyen, et al. Principles of rna processing from analysis of enhanced clip maps for 150 rna binding proteins. Genome biology, 21:1–26, 2020.

45. D. Van Tran, N. Navarin, and A. Sperduti. On filter size in graph convolutional networks. arXiv preprint arXiv:1811.10435, 2018.

46. Z. Wu, S. Pan, F. Chen, G. Long, C. Zhang, and P. S. Yu. A comprehensive survey on graph neural networks. arXiv preprint arXiv:1901.00596, 2019.

47. B. J. Zarnegar, R. A. Flynn, Y. Shen, B. T. Do, H. Y. Chang, and P. A. Khavari. irclip platform for efficient characterization of protein–rna interactions. Nature methods, 13(6):489, 2016.

48. M. Zhang, Z. Cui, M. Neumann, and Y. Chen. An end-to-end deep learning architecture for graph classification. In Thirty-Second AAAI Conference on Artificial Intelligence, 2018.

